# The Genetic Architecture of Quantitative Traits Cannot Be Inferred From Variance Component Analysis

**DOI:** 10.1101/041434

**Authors:** Wen Huang, Trudy F.C. Mackay

## Abstract

Classical quantitative genetic analyses estimate additive and non-additive genetic and environmental components of variance from phenotypes of related individuals. The genetic variance components are defined in terms of genotypic values reflecting underlying genetic architecture (additive, dominance and epistatic genotypic effects) and allele frequencies. However, the dependency of the definition of genetic variance components on the underlying genetic models is not often appreciated. Here, we show how the partitioning of additive and non-additive genetic variation is affected by the genetic models and parameterization of allelic effects. We show that arbitrarily defined variance components often capture a substantial fraction of total genetic variation regardless of the underlying genetic architecture in simulated and real data. Therefore, variance component analysis cannot be used to infer genetic architecture of quantitative traits. The genetic basis of quantitative trait variation in a natural population can only be defined empirically using high resolution mapping methods followed by detailed characterization of QTL effects.

## Introduction

Classical quantitative genetic analysis begins with the premise that the amount of phenotypic variation of quantitative traits in a natural population is determined by the genetic variation of the trait and environmental variation that is random in nature (Fisher 1918; Falconer and Mackay 1996). Genetic variation of quantitative traits can be further partitioned into the additive variance (*V*_*A*_)], dominance variance (*V*_*D*_), and inter-locus interaction (epistatic) variance (*V*_*I*_) (Falconer and Mackay 1996). All of the genetic variance components can be defined in terms of an underlying genetic model with additive, dominance and epistatic genotypic effects and allele frequencies (Table 1) (Fisher 1918). These quantitative genetic variance components are key parameters in theoretical treatments of quantitative trait variation. The additive genetic variance *V*_*A*_ is of particular importance because it defines the level of narrow sense heritability (*h*^2^), which determines the rate of response to natural or artificial selection (Lush 1943).

**Table 1.** **Notations and definitions of variance components in this study**.

| Notation | Variance component | Genotype coding |
| --- | --- | --- |
| $V_A$ | $2pq[a + d(p - q)]^2$ | $x_A \in \{0, 1, 2\}$ |
| $V_D$ | $(2pqd)^2$ | $x_D \in \{0, 2p, 2(p - q)\}$ |
| $V'_D$ | $\frac{4pq^2}{1 + q}(a + dq)^2$ | $x'_D \in \{0, 2, 2\}$ |
| $V'_A$ | $\frac{2p^2q}{1 + q}(a - d)^2$ | $x'_A \in \{0, \frac{1 - q}{1 + q}, \frac{-2q}{1 + q}\}$ |
| $V''_{AA}$ | computed numerically | $x''_{AA} \in (x_{A,1} - 1)(x_{A,2} - 1)$ |

The relative importance of *V*_*A*_, *V*_*D*_, and *V*_*I*_ and especially the additive, dominant, and epistatic gene actions they imply is controversial. Although genetic mapping studies frequently detect non-additive intra-and inter-locus interactions, variance component estimates nearly unanimously find that *V*_*A*_ is the majority of the genetic variance, and dwarfs the contribution of *V*_*D*_ and *V*_*I*_ (Hill *et al*. 2008; Bloom *et al*. 2013; Mäki-Tanila and Hill 2014; Zhu *et al*. 2015). These studies are the basis for the argument that *V*_*A*_ is the overwhelming determinant of genetic variation. Therefore, it is often stated that non-additive variance - and implicitly non-additive gene action - plays only a minor role in the genetic architecture of quantitative traits because non-additive gene action mostly contributes to *V*_*A*_; and because the magnitude of *V*_*D*_ and *V*_*I*_ from dominance and epistasis, respectively is small (Hill *et al*. 2008; Mäki-Tanila and Hill 2014).

However, there are many and potentially an infinite number of possible ways other than *V*_*A*_, *V*_*D*_, and *V*_*I*_ to partition genetic variance, all of which are dependent on allele frequencies in natural populations (Zeng *et al*. 2005). The implicit equivalence of just one of the ways, *V*_*A*_, *V*_*D*_, and *V*_*I*_ with additive, dominant, and epistatic gene action respectively is problematic. Because *V*_*A*_ seeks to maximize the variance it explains, unless genotype frequencies meet specific conditions determined by the genetic models, all types of gene actions can contribute to *V*_*A*_ (Hill *et al*. 2008), making it impossible to use variance components to quantify the contributions of them. However, with a few exceptions, for example, by distinguishing statistical and physiological epistasis (Cheverud and Routman 1995; Alvarez-Castro and Carlborg 2007), there has been a general lack of effort to distinguish variance components from the underlying gene actions that contribute to them, which has caused great confusion. Here, we show that the partitioning of genetic variance components is dependent on the parameterization of allelic effects, and that arbitrarily defined variance components have similar variance explaining ability as *V*_*A*_, regardless of genetic models. We argue that the classical definition of *V*_*A*_ is often inappropriate in the context of explaining within-generation variance, and that partitioning of genetic variance in general provides no information regarding the underlying genetic architecture of quantitative traits.

## Results

### Additive genetic variance *V*_*A*_ is a major determinant of total genetic variance

Following conventional notation, we arbitrarily assign the genotypic value of the three possible genotypes aa, Aa, and AA at a single bi-allelic locus as - *a, d*, and +*a* respectively (Falconer and Mackay 1996). Additive and dominant gene actions (or “genetic model”) have a clear meaning with this parameterization. An “additive” genetic model refers to the situation in which *d* = 0, and hence there is a completely linear relationship between the genotypic value and the number of copies of A alleles. A “dominant” genetic model is when *d* = ±*a*, or when the genotypic value is solely determined by the presence of the dominant allele. *V*_*A*_ (see Table 1 for this and other notations and definitions used throughout this study) accounts for the entirety of genetic variation when the true genetic model is an additive model (Figure 1a). *V*_*A*_ also explains the majority of genetic variation under the dominant genetic model unless the dominant allele is at high frequency (Figure 1b). Extending this single-locus model to two unlinked loci, it can be shown that *V*_*A*_ captures the majority of overall genetic variance unless both loci are frequent under a two-locus “additive by additive” genetic model (Figure 1c,d). These simple results have been previously shown by many authors (Falconer and Mackay 1996; Hill *et al*. 2008; Mackay 2014) but are reproduced here to set the stage for the following results.

**Figure 1.**
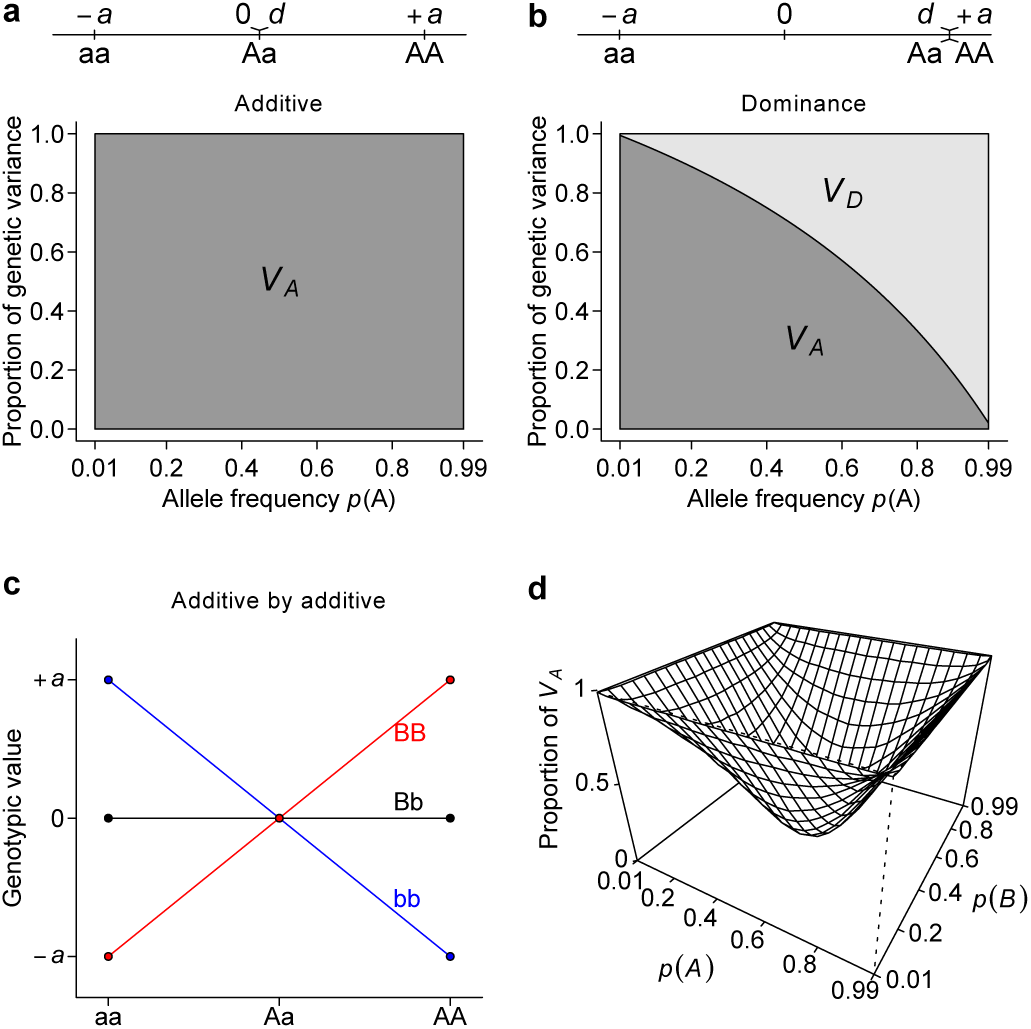
Additive genetic variance *V*_*A*_ is a major determinant of total genetic variance. Under additive (**a**), dominant (**b**), or additive by additive (**c, d**) models, the proportion of total genetic variance explained by the additive genetic variance *V*_*A*_ and dominance genetic variance *V*_*D*_ are estimated either analytically (**a, b**) or numerically by simulation (**d**).

### Alternative parameterizations also capture the majority of genetic variance

The discrepancy between the well-defined genetic models and the literal implication of the term additive genetic variation (*V*_*A*_), *i.e*., apparently non-additive genetic models (dominant or additive by additive) produce significant additive variance, is confusing and counterintuitive. Realizing that *V*_*A*_ is a population property and relies on a specific parameterization of genotypes, we derive alternative parameterizations and quantify the variance explained by them (Table 1). Using a single-locus parameterization in which the heterozygotes and the homozygotes for the dominant allele are coded identically, we define an alternative dominance variance 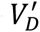 (Table 1, the prime symbol is used to distinguish this variance from the conventional dominance variance *V*_*D*_), which is found to capture the entire genetic variance when the true genetic model is a dominant model (Figure 2a). Importantly, similar to the phenomenon in the classical parameterization in which *V*_*A*_ absorbs the majority of the genetic variance, under this parameterization 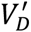 can explain a substantial amount of genetic variance even when the true genetic model is additive (Figure 2b). Furthermore, an alternative two-locus parameterization (see Methods) allows a newly defined 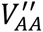 variance component (Table 1) to explain the entire genetic variance with an additive by additive genetic model (Figure 2c) while still capturing a majority of genetic variance under most circumstances when the genetic model is purely additive (Figure 2d).

**Figure 2.**
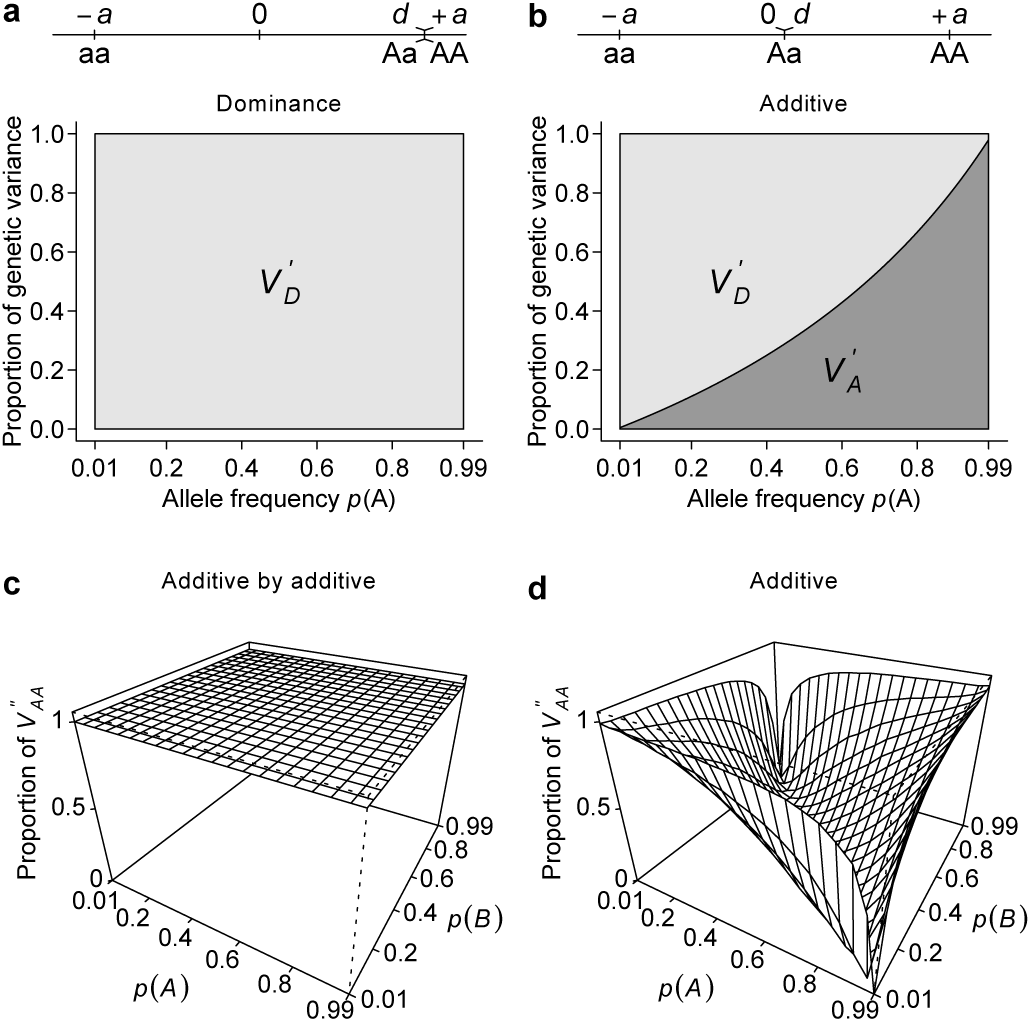
Alternative parameterizations capture the majority of genetic variance. Using an alternative parameterization that emphasizes dominant gene action, a newly defined dominance variance 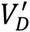 and additive deviation variance 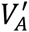 are estimated analytically under dominant (**a**) and additive (**b**) models. Using an alternative parameterization that emphasizes additive by additive gene action, a newly defined interaction variance 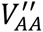 is estimated numerically under additive by additive (**c**) and additive (**d**) models.

### Conventional and alternative parameterizations capture the majority of polygenic genetic variance

To extend the single- and two-locus results to polygenic genetic models, we simulated genotypes and phenotypes based on pre-defined genetic architecture and broad sense heritability (*H*^2^), and used mixed models to partition phenotypic variance under the classical and alternative parameterizations described above. As expected, when the genetic parameterizations and the resulting genetic covariance matrices match the true genetic models, the estimated variances fully explain the total genetic variances (Figure 3). Intriguingly, similar to the single-and two-locus models, all genetic parameterizations are able to capture a major (almost always > 40%) fraction of total genetic variances regardless of the true genetic architecture (Figure 3). Among the three parameterizations, the classical definition of *V*_*A*_ appears to explain the most genetic variance when the genetic model does not match its parameterization. This is likely because the genotypic coding under the conventional additive parameterization is insensitive to the sign of the allelic effects; while the dominance parameterization requires prior knowledge of the dominant allele, and the additive by additive parameterization requires prior knowledge of the interacting pairs. Nonetheless, it is remarkable that even with random assignment of the dominant allele or random pairing of loci and obvious mischaracterization of the genetic model, 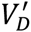 and 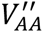 are able to explain the majority of genetic variance when the genetic architecture is additive within and between loci (Figure 3b,c).

**Figure 3.**
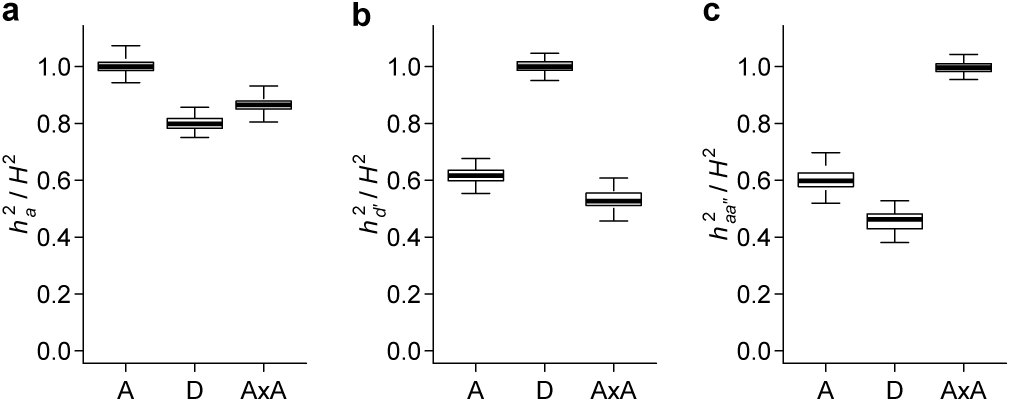
Conventional and alternative parameterizations capture the majority of polygenic genetic variance. Simulation is used to generate data sets with the additive (A), dominant (D), and additive by additive (AxA) genetic models and *V*_*A*_, 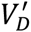, and 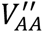 are estimated using linear mixed models. The results are presented as the proportion of heritability explained by the genetic variance component; 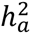 corresponds to *V*_*A*_, 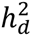, to 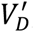, and 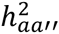 to 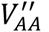.

### *V*_*A*_, 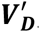, and 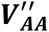 explain a large fraction of phenotypic variance for human height

It has been previously shown that *V*_*A*_ accounts for a large fraction of phenotypic variance in human height using a genetic covariance matrix computed from genome-wide SNP data under the conventional parameterization (Yang *et al*. 2010). Based on our above results, we necessarily expect this result regardless of genetic architecture and indeed recapitulated it using genotype and height data for individuals from the GENEVA project (Figure 4). We then asked if 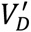 and 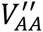 under alternative parameterizations can do the same as in simulated data. Remarkably, and under the naive assumptions that minor alleles are recessive and randomly pairing interacting SNPs, both 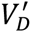 and 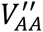 can explain a substantial fraction of phenotypic variance, with even larger point estimates than *V*_*A*_ (Figure 4).

**Figure 4.**
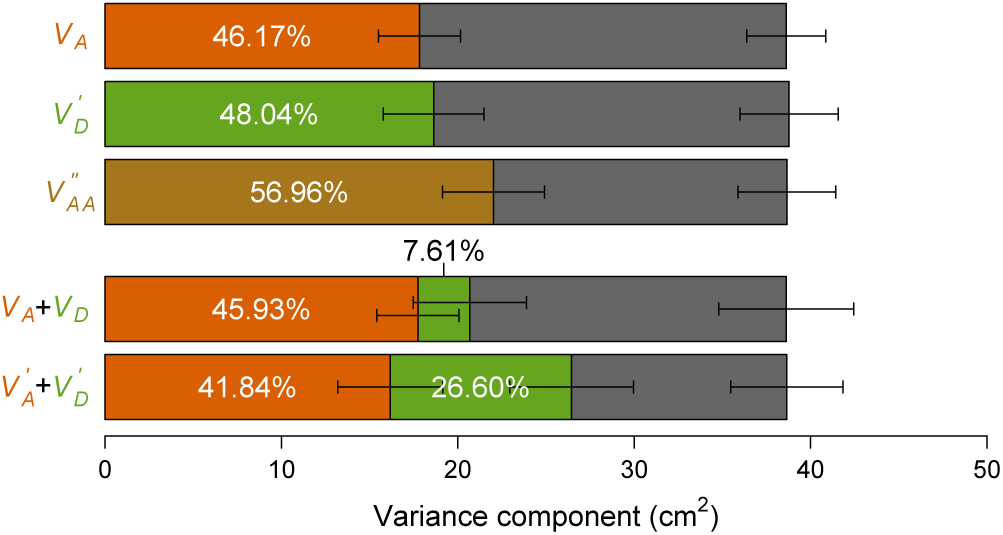
Variance component analyses of human height data. Phenotypic variation of height (in cm) observed in the GENEVA study is partitioned into genetic variance components as indicated (color-coded bars) and environmental variance (***V***_***e***_, grey bar). The colors of bars correspond to the colors of the text indicating the variance components. Error bars indicate standard errors of the variance component estimates provided by GCTA. Proportions of the components are also indicated.

A recent study reported a major contribution of *V*_*A*_ and a minor contribution of *V*_*D*_ for a number of quantitative traits, using the classical parameterization for the additive genetic variance to estimate *V*_*A*_, and a frequency-dependent parameterization (Table 1) orthogonal to the additive genetic value to estimate *V*_*D*_ (Zhu *et al*. 2015). This observation led to the conclusion that dominance variation contributes little to quantitative trait variation. Using the same method, we also observed similar relative contributions of *V*_*A*_ and *V*_*D*_ for human height in the GENEVA data (Figure 4). However, this is only one of the many possible ways of partitioning variance. Using our alternatively defined parameterizations and a similar frequency-dependent parameterization orthogonal to the dominance genetic value (Table 1), we find a much more substantial contribution of dominance variance, *i.e*., 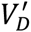 (Figure 4), further suggesting that partitioning of genetic variance is dependent on parameterization.

## Discussion

In animal breeding, *V*_*A*_ and *h*^2^ estimated from pedigree data inform us about the proportion of phenotypic variation that is “breedable”, which is precisely how these genetic parameters were initially defined and used. On the other hand, while broad sense heritability (*H*^2^) is the true measure of the contribution of all sources of genetic variation, it is impossible to estimate in most natural populations, including humans, because no replicated measures can be made on genetically identical individuals, and therefore *h*^2^ provides a lower bound of *H*^2^. Our results and those of many others clearly indicate that it is a reasonable lower bound because in most cases *h*^2^ is close to *H*^2^.

While *V*_*A*_ is certainly useful to give an indication of the magnitude of *H*^2^, our results show that it is not the only one. Alternative parameterizations and the variance they explain, 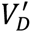 and 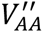, do similarly well in most cases in simulated as well as real data. The emphasis on *V*_*A*_ in animal breeding is sensible because *V*_*A*_ but not 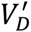 or 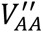 determines the potential of genetic progress (Lush 1943). However, there is little reason to take this *V*_*A*_ centric perspective when explaining within-generation genetic variation. In natural populations, whether one of these variance components is more useful than another depends on the true underlying genetic models. Though it is impossible to determine a clear winner, it is obvious that *V*_*A*_ fully explains genetic variation under an additive model, 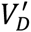 under a dominant model, and 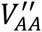 under an additive by additive epistatic model. Therefore it seems only appropriate to define *V*_*A*_ as the additive variance when the genetic model is additive, but to define 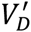 rather than *V*_*D*_ as the dominance variance when the genetic model is dominant, and to define 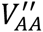 as the epistatic variance when the genetic model is entirely additive by additive. However, these definitions are only valid under very specific and strict circumstances, limiting their use.

Because of such strong dependency of genetic variance components on genetic models, the literal meaning implied by conclusions such as “the majority of genetic variance is due to additive variance” is meaningless; the same could be said for 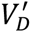 and 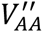. The definition of *V*_*A*_ as “additive” variance is thus an inappropriate one because it reflects the presumed genetic model and genetic parameterization rather than the true genetic model. The relative importance of estimated genetic variance components is also meaningless because it changes with the parameterizations. Therefore, genetic variance components do not inform us about the underlying genetic architecture. This is true for *V*_*A*_, 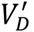, and 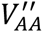 and independent of the parameterization used.

The ability of arbitrarily defined parameterizations to capture the majority of genetic variance shares analogy with the ability of the type I sum of squares to explain variance that is not always attributable to the experimental factor when the experimental design is not orthogonal. In genetic studies, an orthogonal design is not always achievable and impossible in natural populations. While orthogonal parameterizations and partitioning of variance are possible subsequently (Zeng *et al*. 2005) (e.g. the portioning of *V*_*A*_ and *V*_*D*_ are based on orthogonal parameterizations, so are 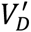 and 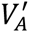), they are fundamentally different from a hypothetical designed orthogonal experiment, in which different gene actions can be applied independently as treatments. Importantly, the additive and dominant gene actions as commonly defined (Figure 1a,b), are two intrinsically inseparable terms and not independent, with a strictly additive model sometimes referred to as co-dominance. This is the root of the confusing convolution of different variance components, especially when not clearly defined.

In the modern era with genome-wide genetic polymorphism data, the segregation of quantitative trait loci (QTLs) can be tracked by DNA markers in linkage or linkage disequilibrium with the true QTLs; and, in rare cases, the true QTLs themselves. Therefore it is possible to compare the observed genetic models with the presumed models and to define genetic architecture once the participating players have been identified. Genome wide association studies (GWAS) have become the most common implementation of marker based approaches. However, QTLs identified by GWAS fail to explain the majority of genetic variation additively, a phenomenon referred to as “missing heritability” (Manolio *et al*. 2009). Simultaneously using all markers additively (Yang *et al*. 2010), as well as using our newly defined parameterizations assuming alternative genetic models (Figure 3,4), can explain a large fraction of genetic variation. Although this holistic method cannot identify QTLs and offers no information on the underlying genetic models, it suggests that the heritability is just missing but does not vanish. Strategies to find the heritability include increasing sample size to find QTLs of smaller size, using sequencing to find QTLs of rarer frequencies, and testing for non-additive effects. Unfortunately, the effort to model non-additive effects is limited because of its requirement for much larger sample size and the predominant assumption that non-additive effects are unimportant. Our results suggest that this assumption is unfounded. Undoubtedly, it is experimentally and statistically challenging to find all QTLs, even more so to search for cryptic genetic models other than additivity and dominance within a single locus, including two way or higher order interactions, and other cryptic genetic models. Like the single locus models, these more complex genetic models may not ultimately explain the missing heritability. Nonetheless, it is clear that the current standard single-locus approach is not adequate precisely because of the missing heritability phenomenon itself.

## Acknowledgement

This work was supported by NIH grants R01 GM45146, R01 AA016560 and R01 AG043490 to TFCM.

## Methods

### Least squares regression interpretation of *V*_*A*_

Consider a single biallelic locus in a diploid genome with alleles A and a, each with frequency p and *q* (*p* + *q* = 1); and assign genotypic values *y* = -*a,d*, and +*a* to genotypes aa, Aa, and AA respectively. The average effects of A and a are then *qα* and -*pα* respectively, where *α* = *a* + *d*(*q* – *p*) is the allele substitution effect and measures the change in phenotype in an individual if an allele a is substituted with A(Falconer and Mackay 1996). The breeding value, defined as the expected genotypic value of the progeny an individual produces, is the sum of average allelic effects each diploid individual carries, and is - 2*pα*, *qα*; – *pα*, and 2*qα* for aa, Aa, and AA respectively. With only one locus, the total genetic variation in a randomly mating (thus in Hardy-Weinberg equilibrium) population can be partitioned into two orthogonal components, the additive genetic variance *V*_*A*_, which is defined as the variance due to breeding values, 2*pqα*^2^, and the dominance genetic variance *V*_*D*_ = (2*pqd*)^2^ (Table 1)(Falconer and Mackay 1996).

Alternatively, we can define a random variable *x*_*A*_ as:

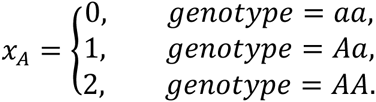

This parameterization has the convenient interpretation that *x*_*A*_ is equal to the number of A alleles. It is easy to show that the allele substitution effect *α* as defined above is the slope of the least squares regression of genotypic value *y* on *x*_*A*_ in an idealized population with random mating (Figure S1). The additive genetic variance is then *V*_*A*_ = *Var*(*ŷ*) = *Var*(*αx*_*A*_) = *α*^2^*Var*(*x*_*A*_) = 2*pqα*^2^ and the dominance genetic variance *V*_*D*_ is the residual variance. It is easy to see that the least squares solution for this regression seeks to maximize *V*_*A*_ and minimize *V*_*D*_. This least squares interpretation is not new and dates back to the early days of quantitative genetics(Fisher 1918).

By extension of this least squares regression interpretation of genetic variation, if we arbitrarily define any one random variable *x* or more than one of them and fit a linear model of form *y* = *βx* + *ϵ*, we can partition genetic variance due to the assumed genetic model *Vari*(*ŷ*) = *Var*(*βx*) and residual variance *Var*(*ϵ*).

### Derivation of dominance variance 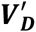 using least squares regression

Now we illustrate the idea of using least squares regression to partition genetic variance due to dominant gene action and the remaining genetic variance. We define the random variable 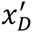 as:

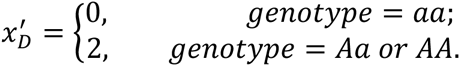

The least square solution for the linear model 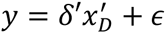 can be easily found to be 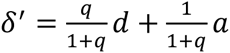 (Figure S2). Therefore the variance due to 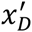 is 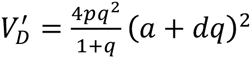. The residuals from this regression are 0,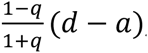, and 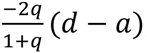, for genotypes aa, AA, and AA respectively. Similar to *V*_*A*_ and *V*_*D*_, we define the residual variance as an “additive deviation” variance 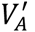, which can be found to be 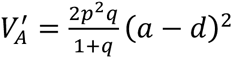.

### Finding 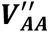 numerically

Extending the least squares regression interpretation of genetic variance to any arbitrary random variable x and finding the solution is not always easy. However, it is computationally trivial to find. For example, to numerically estimate the additive by additive variance 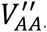, we define 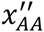 as follows for two independently segregating loci with alleles A/a, and B/b respectively:

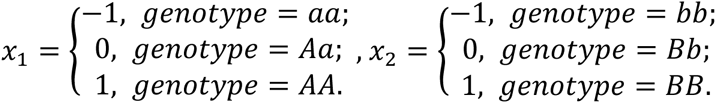

Then, 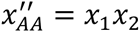. We randomly draw 100,000 individuals with the specific genotypes according to pre-defined allele frequencies and assign genotypic values with pre-defined genetic models. The slopes 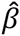 can be easily found by numerically regressing *y* onto *x*. The proportion of genetic variation explained by this parameterization is then just the *R*^2^ of the regression.

### Mixed model analysis of simulated and real data

To extend the single-and two-locus models to polygenic models, we used mixed model analysis to partition phenotypic variation in simulated and real data. To simulate phenotypic data with pre-defined genetic models, we first drew from the U-shaped distribution(Hill *et al*. 2008) 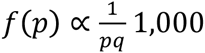 realizations, which took possible values of 0.01, 0.02,…, 0.99. Genotypes for these *p* = 1,000 loci were randomly assigned according to their Hardy-Weinberg frequencies to *n* = 5,000 individuals. Genetic values were then assigned to the 5,000 individuals using this general formula **g = Xβ**. Each of the columns of the *n*×*p* matrix **X** was coded by the additive parameterization *x*_*A*_ as defined above for the additive genetic model, 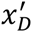 for the dominance genetic model. Similarly for the additive by additive genetic model, 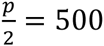 pairs of loci were parameterized as defined above using 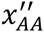. The vector **β** was drawn from standard normal distribution. The phenotypic value *y* for each individual was then simulated by adding random noise such that **Y = g + ϵ**, where **ϵ** was normally distributed with zero mean and variance equal to 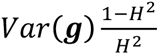. *H*^2^ was the broad sense heritability and was always set to 0.5.

We standardized columns of **X** and computed the covariance matrix as **XX^T^**, which was further scaled by the mean of its diagonal values. A linear mixed model **Y** = *μ***1** + **Zu** + **ϵ** was fitted to the data, where μ was the population mean, **Z** was the incidence matrix and in all cases in this study the identity matrix, ***u*** was a random effect with variance covariance matrix **G***σ*^2^, where **G** was simply the scaled **XX^T^** above and *σ*^2^ was the part of genetic variance due to the specific parameterization. We fitted this model using the GCTA software(Yang *et al*. 2011) with REML and performed simulations 100 times. We defined the heritability explained by *σ*^2^ as *h*^2^ /*H*^2^, where 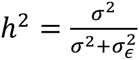, and *H*^2^ was the simulated broad sense heritability.

To analyze real data where the genetic architecture cannot be known *a priori*, we downloaded genotype and phenotype data from dbGaP for the GENEVA Genes and Environment Initiatives in Type 2 Diabetes study (phs000091.v2.p1). We pruned the data set to contain 5,497 unrelated (nominal genetic relationship as calculated by GCTA < 0.05) individuals with European ancestry based on both self-reported ethnicity and principal component analysis. We then computed genetic covariance matrices as defined above using autosomal SNPs and partitioned phenotypic variance using GCTA where sex was fitted as a fixed effect in the model. We used the parameterization (Table 1) as defined in a recent study(Zhu *et al*. 2015) to partition phenotypic variance into *V*_*A*_, *V*_*D*_, and *V*_*e*_. We also partitioned phenotypic variance into 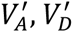, and *V*_*e*_, where 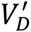 was defined as above and 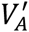 was estimated by defining a new variable 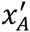 (Table 1), where

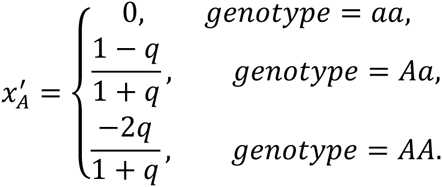

## Figure Legends

**Figure S1.**
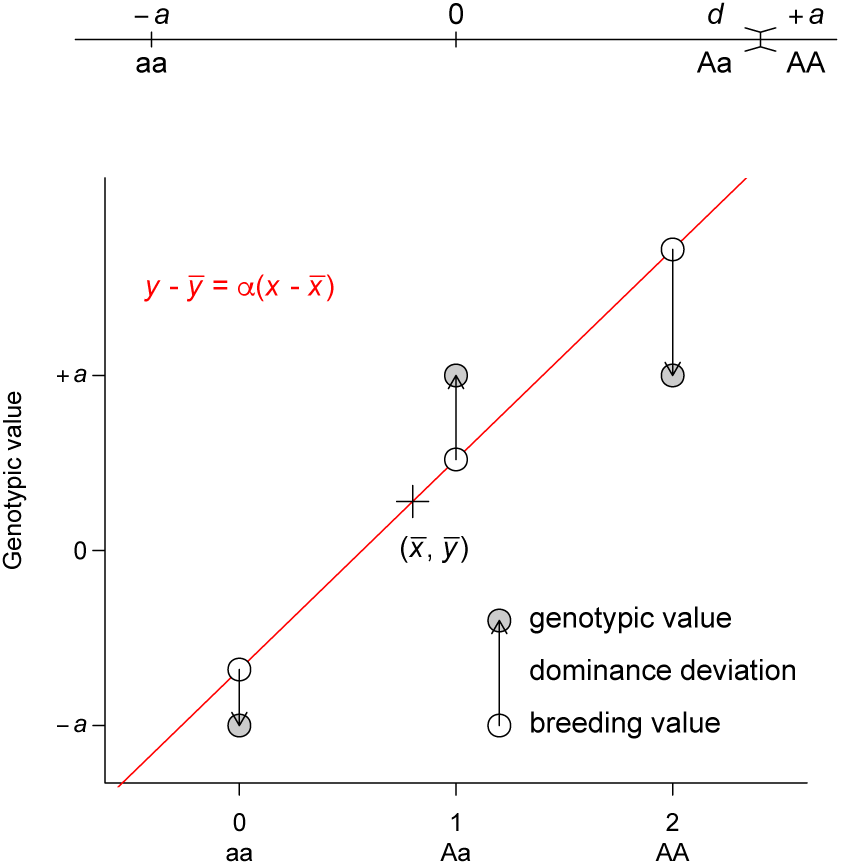
Least squares regression interpretation of *V*_*A*_. This representation is adapted from Fig. 7.2 of Reference 2. Grey circles indicate the genotypic value of each genotype, which is coded as 0, 1, 2 for aa, Aa, and AA respectively. A regression line (red line) is fitted to the data, on which the fitted values are indicated by white circles. The fitted line must pass through the center of the data, as indicated by the cross. The fitted values are equivalent to breeding values. The arrows between the breeding values and the genotypic values are the dominance deviations, which are the same as residuals of the regression. Note that the data points are weighted by their frequencies in the population. A dominance model is used so that the dominance deviation can be illustrated.

**Figure S2.**
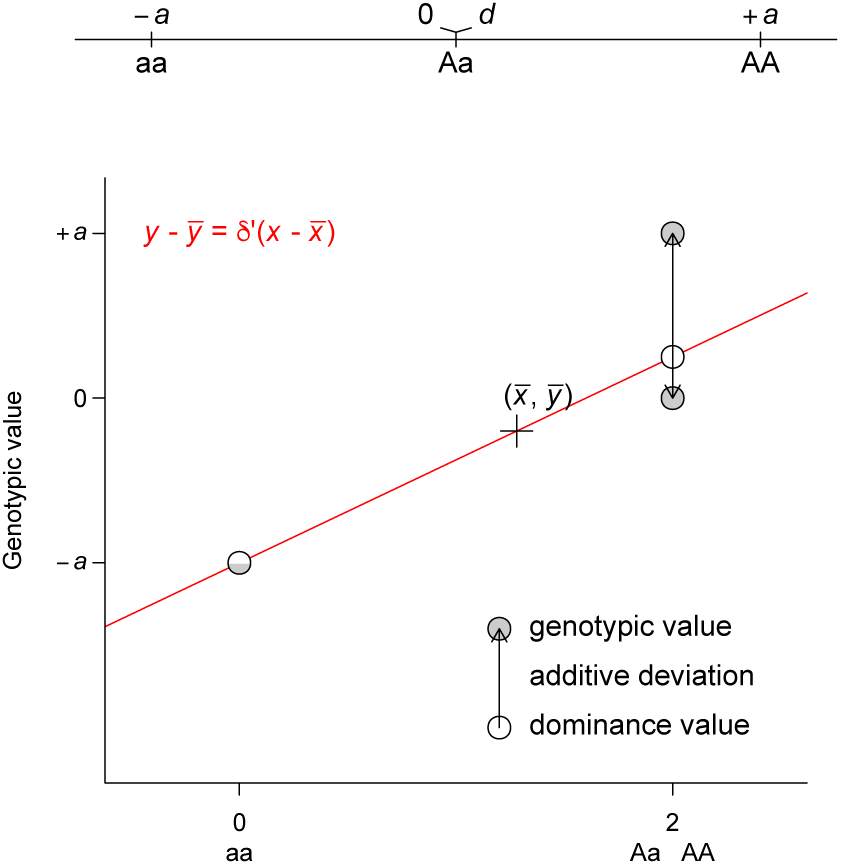
Least squares regression interpretation of 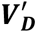. Grey circles indicate the genotypic value of each genotype, which is coded as 0, 2, 2 for aa, Aa, and AA respectively. A regression line (red line) is fitted to the data, on which the fitted values are indicated by white circles. The fitted line must pass through the center of the data, as indicated by the cross. The fitted line must also pass through the circle (half grey and half white to indicate the overlap of the genotypic and fitted values) denoting genotype aa. The fitted values are equivalent to dominance values as defined in this parameterization. The arrows between the dominance values and the genotypic values are the residuals of the regression, which we define as “additive deviation”, therefore the residual variance is 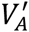. Note that the data points are weighted by their frequencies in the population. An additive model is used so that the additive deviation can be illustrated.

